# Lineage-specific diversification in the usage of D-glutamate and D-aspartate in early-branching metazoans

**DOI:** 10.1101/2020.04.26.036582

**Authors:** Leonid L. Moroz, Dosung Sohn, Daria Y. Romanova, Andrea B. Kohn

**Affiliations:** Whitney Laboratory for Marine Bioscience, University of Florida, St. Augustine, FL, 32080, USA; Institute of Higher Nervous Activity and Neurophysiology, Moscow 117485, Russia; Departments of Neuroscience and McKnight Brain Institute, University of Florida, Gainesville, FL, 32610, USA

**Keywords:** Placozoa, Ctenophora, Porifera, Cnidaria, Capillary electrophoresis, D-amino acids, Evolution, Neurotransmitters

## Abstract

D-amino acids are unique and essential signaling molecules in neural, hormonal, and immune systems. However, the presence of D-amino acids and their recruitment in early animals is mostly unknown due to limited information about prebilaterian metazoans. Here, we performed the comparative survey of L-/D-aspartate and L-/D-glutamate in representatives of four phyla of basal Metazoa: cnidarians (*Aglantha*); placozoans (*Trichoplax*), sponges (*Sycon*) and ctenophores (*Pleurobrachia, Mnemiopsis, Bolinopsis*, and *Beroe*), which are descendants of ancestral animal lineages distinct from Bilateria. Specifically, we used high-performance capillary electrophoresis for microchemical assays and quantification of the enantiomers. L-glutamate and L-aspartate were abundant analytes in all species studied. However, we showed that the placozoans, cnidarians, and sponges had high micromolar concentrations of D-aspartate, whereas D-glutamate was not detectable. In contrast, we found that in ctenophores, D-glutamate was the dominant enantiomer with no or trace amounts of D-aspartate. This situation illuminates prominent lineage-specific diversifications in the recruitment of D-amino acids and suggests distinct signaling functions of these molecules early in the animal evolution. We also hypothesize that a deep ancestry of such recruitment events might provide some constraints underlying the evolution of neural and other signaling systems in Metazoa.

**Highlights:**

- D-amino acids are essential for intercellular signaling and evolution
- Enantiomers have been quantified in early-branching animals
- Lineage-specific recruitment of D-glutamate could occur in ctenophores
- D-aspartate is one of the primary enantiomers in other metazoans
- Deep ancestry of such events could provide constraints in the evolution of signaling

**Graphical Abstract:** D-amino acids are essential for intercellular signaling. Direct microchemical quantification of enantiomers in representatives of early-branching animals suggests lineage-specific recruitments of D-glutamate and D-aspartate. Deep ancestry of such events might provide some constraints underlying the evolution of neural and other signaling systems in Metazoa.

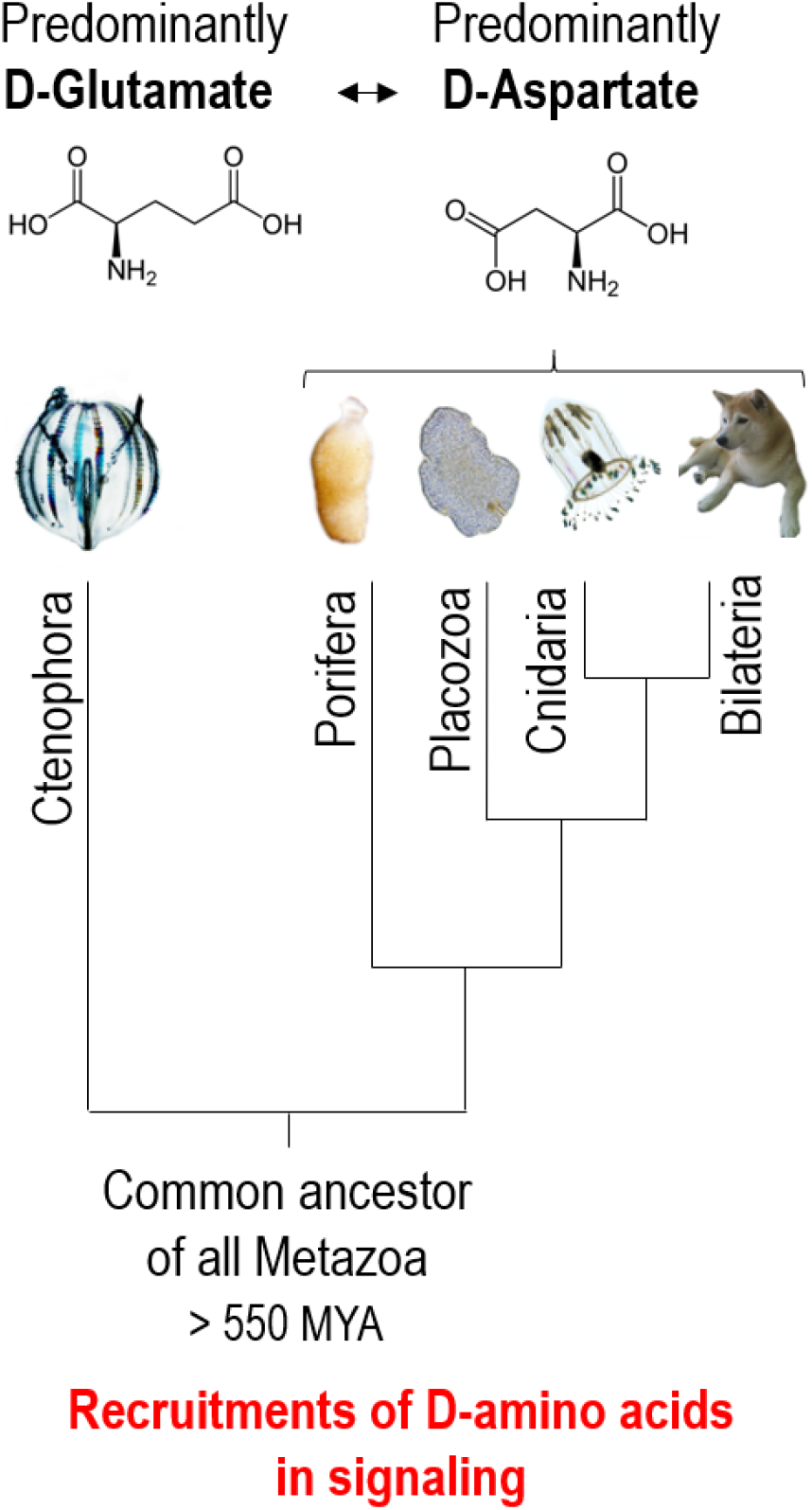

## 1. Introduction

Life is primarily based on L-amino acids, but the growing data from different disciplines stress the recruitments and importance of D-amino acids in various signaling and structural functions at all levels of biological organization [1–3]. With thousands of publications, the chemical biology of D-amino acids is transforming from scattered observations during the 1960-90s [4–6] into a rapidly growing field with many promises for therapeutic and bioengineering [3,7]. For example, since bacteria are known sources of D-amino acids, there are specific enzymes and mechanisms, which recognize these enantiomers as a part of the immune response in mammals [8].

One of the most prominent insights came from analyses of L-glutamate-mediated transmission. D-serine and D-aspartate had received a lot of attention [4,6] due to their relative abundance in the brain [9–11], and involvement in synaptic transmission, including glial signaling [12] and neuromodulation via NMDA receptors [13,14]. Surprisingly, D-glutamate was detected in neurons [15,16], but its functions are less understood; and D-glutamate is not as abundant in the mammalian brain as D-aspartate [17]. D-glutamate was likewise found in *Aplysia* neurons [18,19], again at a significantly smaller level. The reason for an apparent ‘deficiency’ of the usage of D-glutamate vs. D-aspartate in mammals and molluscs (and the majority of invertebrates studied so far) is unknown. Structural or chemical limitations might play a role. Still, there could be some evolutionary constraints in the usage of D-amino acids (including their synthesis and inactivation mechanisms), which requires comparative analyses.

Various D-amino acids were identified only in representatives of several bilaterian lineages such as annelids [5,6], arthropods [4,20–22], echinoderms [23], and hemichordates [24]. The most cell-specific microanalytical data exist for molluscs. The presence, release, and neuromodulatory action of D-aspartate have been demonstrated both in cephalopods [25–27] and gastropods [28–32]. D-glutamate distribution is less studied. D-glutamate was found in the testis of the prawn, *Marsupenaeus japonicus* [20], and in the nematode *Caenorhabditis elegans*, where it is involved in the control of development [33]. In the silkworm, *Bombyx mori* D-glutamate might control muscle contractions [21]. Although it is considered that D-glutamate in animals could enzymatically be produced by an amino acid racemase, similarly to the synthesis of D-aspartate and D-serine [34,35], the existence of specialized glutamate racemase has never been reported in animals. One notable exception is an isoform of the aspartate racemase from the acorn worm (*Saccoglossus*), which can catalyze the formation of both D-aspartate and D-glutamate from their L-forms *in vitro* [24].

In summary, virtually nothing is known about even the usage of D-amino acids early in animal evolution. There are no systematic cross-phyla comparisons of the recruitment of different enantiomers, and comparative data are available for only 7 out of 35 extant phyla. It appears that D-glutamate is less abundant compare to D-aspartate in most bilaterians studied so far. Here, we used high-resolution capillary electrophoresis assays to quantify D-amino acids in representatives of four basal metazoan phyla: Cnidaria, Porifera, Placozoa and Ctenophora. And we found evidence for a remarkable split in the recruitment of D-aspartate and D-glutamate at the base of the animal tree of life.

## 2. Materials and methods

### 2.1. Animals

For comparative microchemical assays, we collected animals from plankton at Friday Harbor Laboratories (University of Washington). The list includes the ctenophores, *Beroe abyssicola, Pleurobrachia bachei, Bolinopsis infundibulum*, and the hydrozoan jellyfish, *Aequorea victoria* (Hydrozoa, Cnidaria). The calcium sponge *Sycon sp*. (class Calcarea, Porifera) was collected at the docks of the Friday Harbor Laboratories. We maintained animals for up to one week in running seawater at ambient temperature before experiments.

*Trichoplax adhaerens* (H1 haplotype), 0.3-2mm in diameter, were maintained in the laboratory culture, and animals were fed on rice grains and algae [36].

We dissected specific body parts of ctenophores under a stereomicroscope and washed samples three times (5 minutes) in artificial seawater. We also washed *Trichoplax* before the use, and each individual was separated for the assay. The samples were stored in a PCR tube containing Milli-Q water at −80°C until use.

### 2.2. Microchemical assays of enantiomers

(L-/D-glutamate, L-/D-aspartate) were performed using high-resolution capillary electrophoresis (CE) with laser-induced fluorescence (LIF) detector. The principle and details of the protocol for amino acid separation assay were reported elsewhere[36–38] with some modifications for chiral analyses, which we summarize below.

The CE, coupled with the ZETALIF detector (Picometrics, France), was used for the assay of amino acids. A helium-cadmium laser (325nm) from Melles Griot, Inc. (Omnichrome^®^ Series56, Carlsbad, CA) was used as the excitation source. Before the photomultiplier tube (PMT), the fluorescence was both wavelengths filtered and spatially filtered using a machined 3-mm pinhole. All instrumentation, counting, and high-voltage CE power supply were controlled using DAx 7.3 software.

All solutions were prepared with ultrapure Milli-Q water (Milli-Q filtration system, Millipore, Bedford, MA) to minimize the presence of impurities. Borate buffer (30mM, pH=9.5) was used for sample preparation. All solutions were filtered using 0.2μm filters. The buffers were degassed by ultrasonication for 10 min to minimize the chance of bubble formation. A 75mM *o*-Phthalaldehyde (OPA)/β-mercaptoethanol (β-ME) stock solution was prepared by dissolving 10mg of OPA in 100μL of methanol and mixing with 1mL of 30mM borate and 10μL of β-ME. Stock solutions (10mM) of amino acids were prepared by dissolving each compound in the borate buffer. OPA and β-ME were stored in a refrigerator; fresh solutions were prepared weekly.

All experiments were conducted using a 75cm length of 50μm I.D. × 360μm O.D. fused silica capillary (Polymicro Technologies, AZ). A 30mM borate/30mM sodium dodecyl sulfate (SDS) electrolyte (adjusted to pH [9 to 10] with NaOH), and 15mM borate and 10mM beta-cyclodextrin electrolyte (adjusted to pH=10.0 with NaOH) was used for a separation buffer of chiral analysis of both glutamate and aspartate (Fig.1).

**Figure 1.**
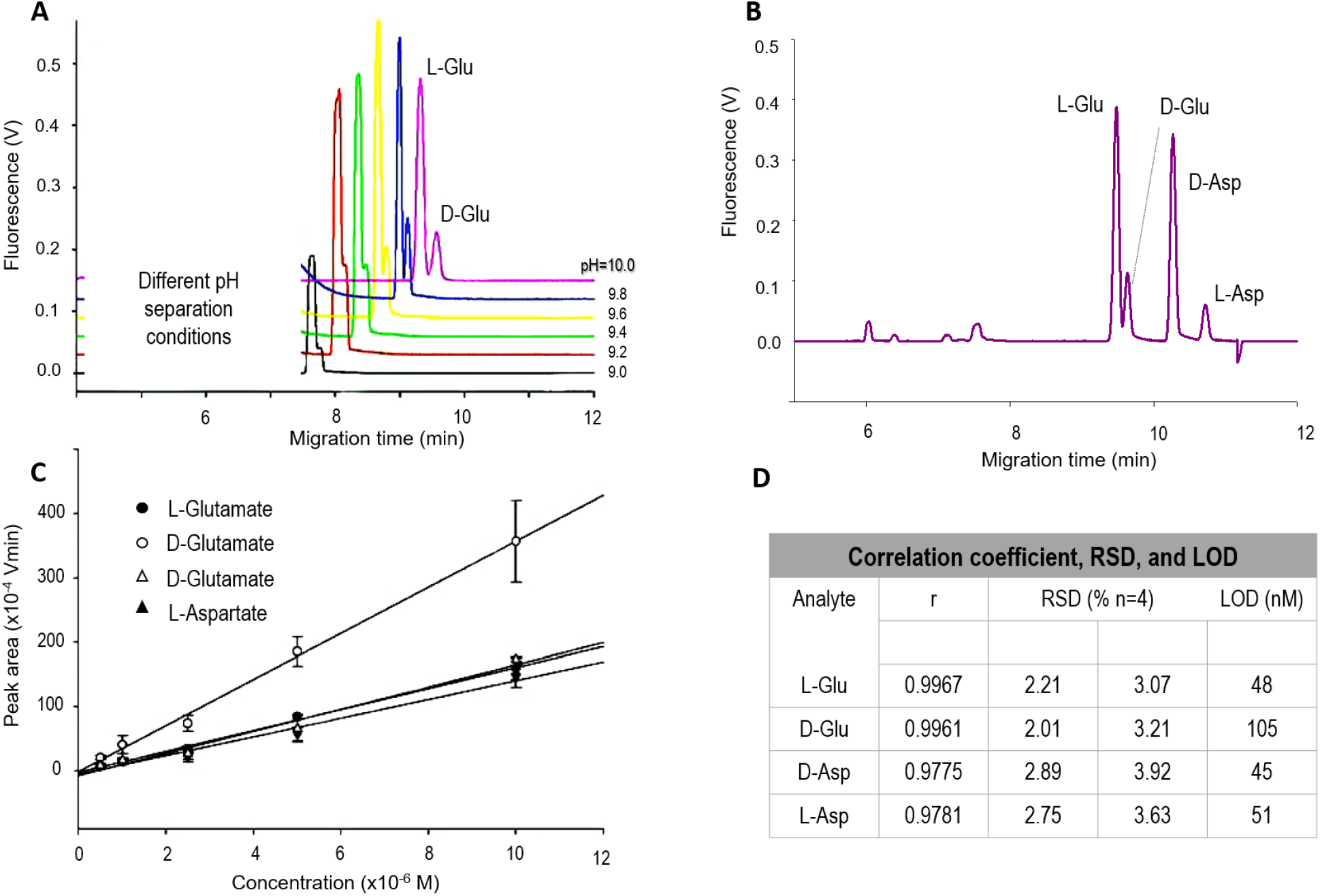
Ultrasensitive Capillary Electrophoresis Assay for Separation of Enantiomers. A and B: Electropherograms of Glutamate and Aspartate enantiomers. A) The separation conditions were pH-dependent (from 9.0 to 10.0) with the optimum at pH=10. B) The electropherograms of 1μM of L-glutamate and D-aspartate, and 100 nM of D-glutamate and L-Aspartate at pH=10.0. C) Standard calibration curves used in the study. D) Correlation coefficients (r), the relative standard deviation (RSDs), and limits of detection (LOD) for the enantiomer separation conditions. Samples were loaded using electrokinetic injection (8 kV for 12 s) and then analyzed under a stable 20 kV voltage at 20°C in 50 μm inner diameter (I.D.) and 360 μm outer diameter (O.D.) capillary with 15 mM borate and 10 mM of beta-cyclodextrin electrolyte (β-CD).

We used the pre-column derivatization method. A 1μL of *o*-Phthalaldehyde (OPA) was placed in a 0.5mL PCR tube. The total volume of a sample, OPA, and internal standard inside the tube was 20μL. For separation steps, the capillary inner-wall was successively washed with 1M NaOH, Milli Q water, and the separation buffer by applying pressure (1900mbar) to the inlet vial. Then the sample was loaded using electrokinetic injection (8kV for 12s). The separation was performed under a stable 20kV voltage at 20°C.

In all CE tests, once an electropherogram was acquired, peaks were assigned based on the electrophoretic mobility of each analyte, and the assignments were confirmed by spiking corresponding standards into the sample. Five-point calibration curves (peak area vs. concentration) of analytes were constructed for quantification using standard solutions, and the dilution factor was required to be considered to determine the initial concentration. We used the 3σ method to determine the limit of detection (LOD). We calculated LOD from standard deviations of blank (n=5) and the calibration slope of low concentration standards. The reproducibility and accuracy of the method were evaluated by calculating the relative standard deviation (RSD) and error values. In order to determine the peak area in the electropherograms, a baseline was constructed and subtracted using a derlim algorithm of DAx software version 7.3 (Van Mierlo Software Consultancy, the Netherlands). Statistical data analysis was performed by Sigma Plot software (SPSS, Inc., Richmond, CA).

All chemicals for buffers were obtained from Sigma-Aldrich, and standard amino acids were purchased from Fluka. We used ultrapure Milli-Q for all solutions and sample preparations.

## 3. Results

### 3.1. Development of a highly sensitive assay for separation of enantiomers using capillary electrophoresis

**Fig. 1** shows the illustrative examples of the separation of the chiral forms of glutamate and aspartate. Since the interactions between beta-cyclodextrin electrolyte (β-CD) and *o*-Phthalaldehyde (OPA)-derivatives are pH-dependent, various pHs were tested to optimize the separation of the glutamate enantiomers (aspartate enantiomers showed significantly lesser dependence on pH). The L and D forms of glutamate closely migrated at pH 9.0, but they were resolved with increasing pH and were entirely separated at pH 10.0 (**Fig. 1A**). The hydroxyl groups of β-CD become negative in a basic condition; with a 20kV separation voltage, the electroosmotic flow is towards the negative electrode, while β-CDs move in the opposite direction[39]. Thus, the more an analyte interacts with the β-CD, the slower the complex moves to the negative electrode. **Fig. 1B** shows that the interaction strength between β-CD and analytes are L-Aspartate>D-Aspartate>D-Glutamate>L-Glutamate.

Calibration curves were constructed by injecting a series of standard mixtures covering the tested concentration range (**Fig. 1C**). Equations were obtained by the least-squares linear regression analysis of the peak area versus analyte concentration. **Fig. 1D** summarizes the results of the determination of reproducibility regarding the accuracy, within-day, and day-to-day precision assays and limits of detection (LOD). The intra-assay precision of the method based on within-day repeatability was performed by replicate injections (n=4), where peak areas were measured. The statistical evaluation provided the relative standard deviations (RSD) at different concentrations. The inter-assay precision (between-day variation) of the method was established by triplicate measurements of each concentration over three different days. The measured concentrations had RSD values <4%.

In summary, we achieved reliable and reproducible assays with nanomolar limits of detection (LOD, **Fig. 1D**).

### 3.2. Detection of endogenous enantiomers in ctenophores, placozoans, sponges, and cnidarians

The current separation protocol, with β-CD as a chiral selector, was highly sensitive. As a result, for the samples obtained from marine animals, we usually performed 1:100 dilutions to be within the range of the detection scheme. In all tests summarized below, the presence of both D- and L-forms was confirmed by spiking of respective analytes for the same sample, immediately after the first assay.

**Figs. 2,3** illustrate examples of these assays for ctenophores and placozoan species, respectively. **Fig. 4** summarizes the quantification of enantiomers in all species studied and their evolutionary relationships.

**Figure 2.**
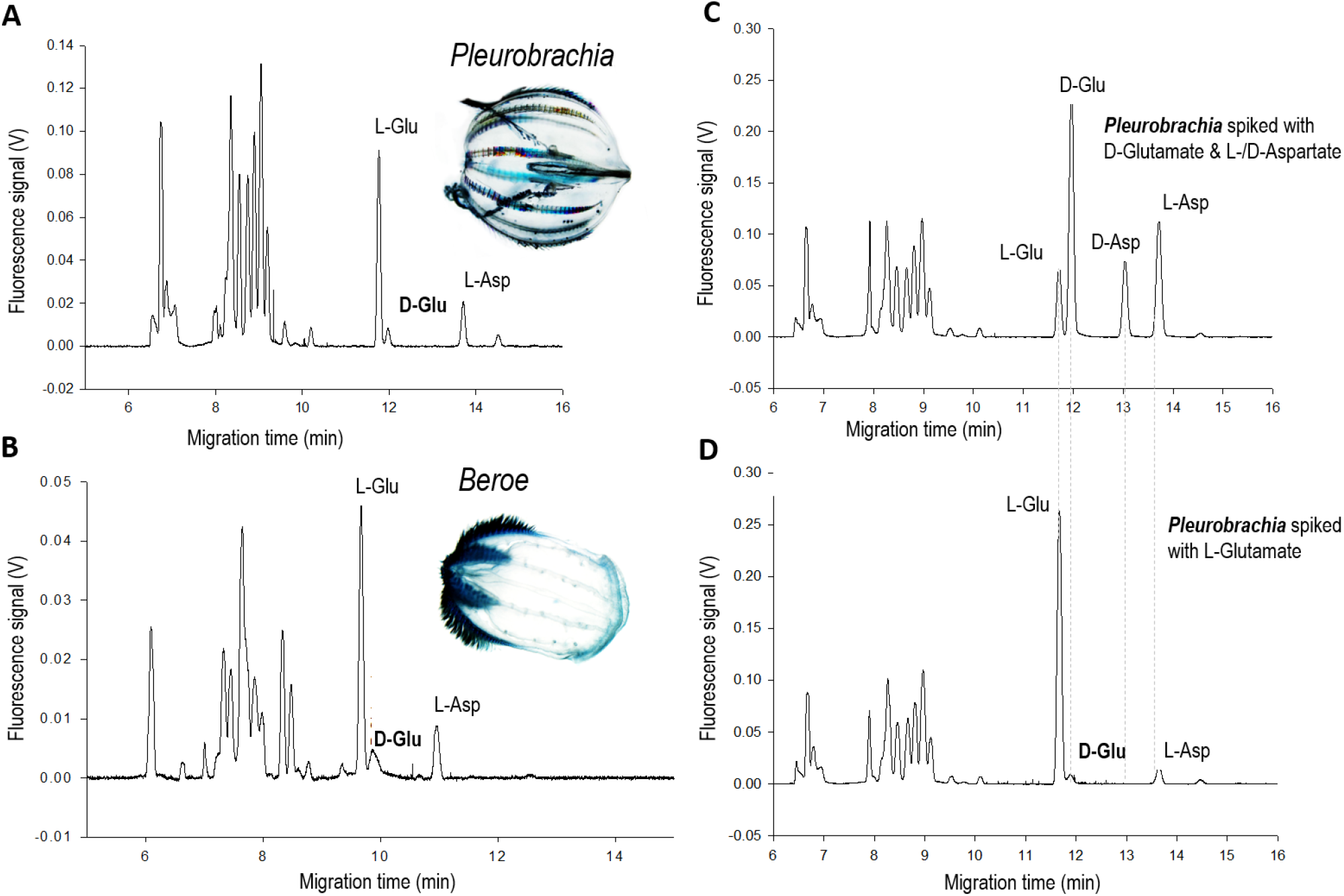
Detection of D-/L-Glutamate and D-/L-aspartate in *Pleurobrachia* and *Beroe* (Ctenophora): D-glutamate as the major endogenous enantiomer. Electropherograms and concentration profile of glutamate and aspartate enantiomers in *Pleurobrachia bachei* (A) and *Beroe abyssicola* (B) – samples represent whole animals. (C) Electropherograms of standards. (C) Electropherograms of *Pleurobrachia* spiked with D-glutamate (1 μM), L-aspartate (1 μM), and D-aspartate (10 μM). (D) Electropherograms of *Pleurobrachia* spiked with L-glutamate (10μM). Note, D-aspartate was not detected in A (*Pleurobrachia*), B (*Beroe*), D (*Pleuronrachia*). Samples were loaded using electrokinetic injection (8 kV for 12s) and then analyzed under a stable 20 kV voltage at 20°C in 50μm inner diameter (i.d.) and 360μm outer diameter (o.d.) capillary with 15 mM borate and 10 mM β-CD, pH=10.0.

**Figure 3.**
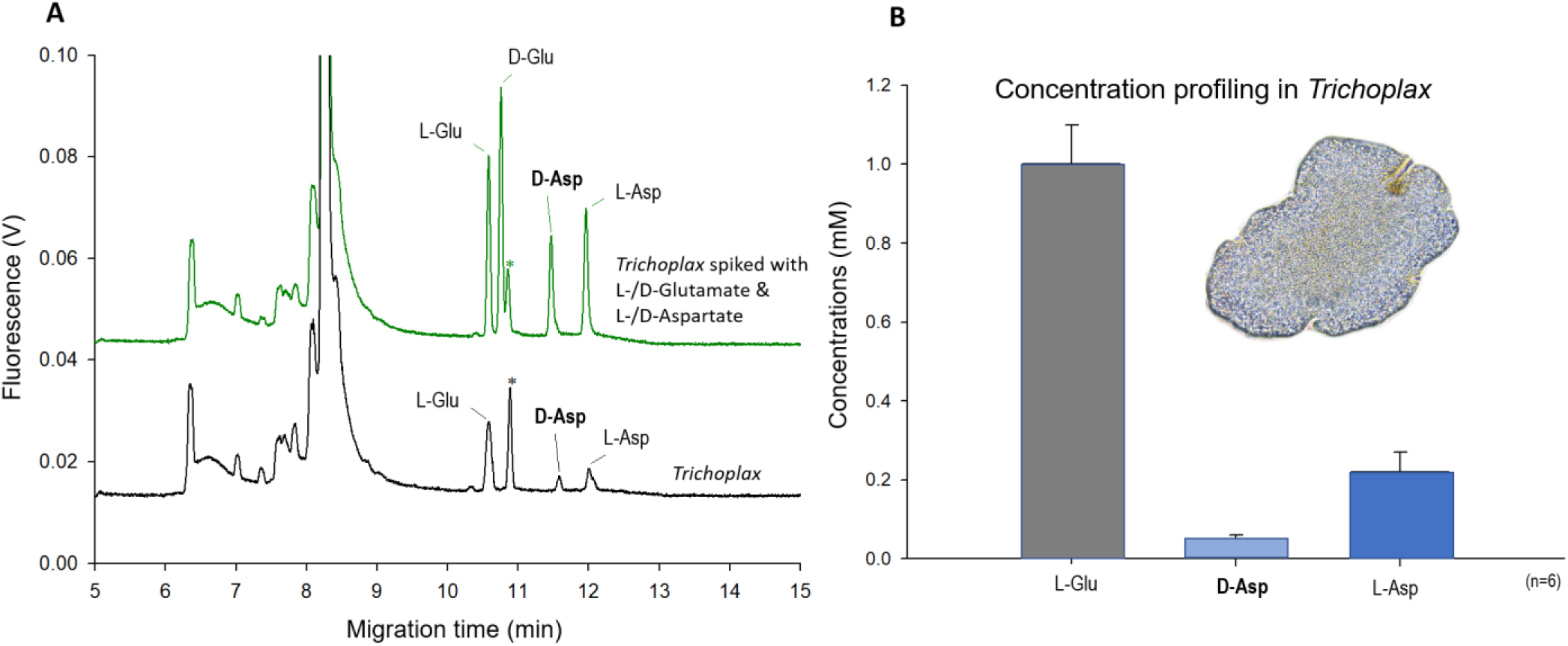
Electropherograms and concentration profile of glutamate and aspartate enantiomers in *Trichoplax adhaerens* (Placozoa): D-aspartate as the major endogenous enantiomer. (A) The green (upper) electropherograms of *Trichoplax* spiked with standards [peaks: L-glutamate (10 μM), D-Glu (1 μM), D-aspartate (10μM), and L-aspartate (1 μM). The black (lower) electropherogram is a *Trichoplax* sample under control conditions. (*) is an unknown peak. Note, D-glutamate was not detected in *Trichoplax*. (B) Concentration profile of glutamate and aspartate in *Trichoplax* (n=4). Samples were loaded using electrokinetic injection (8 kV for 12 s) and then analyzed under a stable 2 0kV voltage at 20°C in 50 μm I.D. and 360 μm O.D. capillary with 15mM borate and 10 mM β-CD, pH=10.0.

Specifically, we profiled four ctenophores species: *Pleurobrachia, Beroe, Mnemiopis*, and *Bolinopsis* (**Figs. 2,4**), which represent distinct phyletic lineages of this earliest-branched animal clade[40]. Considering a relatively complex morphological organization of Ctenophora, we separately analyzed different organs in two species (*Pleurobrachia* and *Beroe*), including the aboral organ – the gravity sensor and an analog of the elementary brain[41,42]. All ctenophore samples produced comparable results with relatively similar concentrations of both L-glutamate and L-aspartate. Surprisingly, we did not detect D-aspartate in ctenophores (**Figs. 2CD, 2S–3S**-supplement). Instead, we identified D-glutamate, which was present at a relatively high level (about 1/10 of the L-glutamate concentrations– **Fig. 4A**). In ctenophores, such apparent “deficit” of D-aspartate and the abundance of D-glutamate contrasts with the profiles of enantiomers in mammals and the majority of studied bilaterians[17,18].

**Figure 4.**
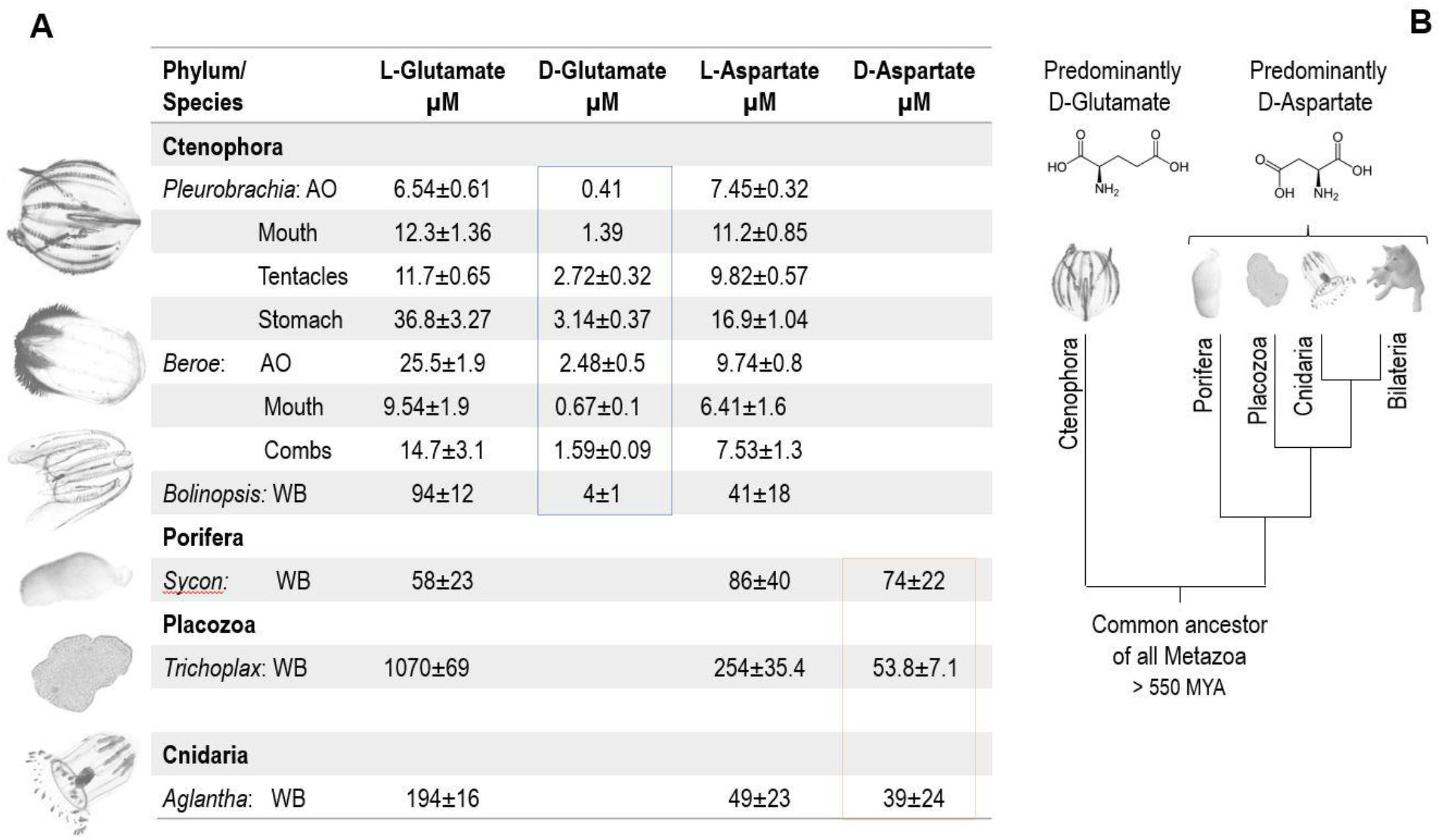
Presence and quantification of D-/L-Glutamate and D-/L-aspartate representative of four phyla of basal Metazoa. (+Supplementary Figs:S2,3S). (A) The table with calculated endogenous concentrations of different enantiomers (μM±SD, n=3-5). AO – aboral organ; WB – whole body. (B) The hypothesis of a possible early split in the recruitments of D-amino acids in different animal lineages. The lineage led to Ctenophores might show the predominant usage of D-glutamate. In contrast, the lineage led to the rest of the animals (including sponges, placozoans, cnidarians, bilaterians) showed the predominant recruitments of D-aspartate[17,18]. The phylogenetic tree is based on the reconstructions summarized elsewhere[40,43,44].

On the other hand, *Trichoplax* had a high level of D-aspartate, L-aspartate, and L-glutamate, but no D-glutamate was detected (**Fig. 3**). Again, the identities of every enantiomer were confirmed by spiking each standard (**Fig. 3A**). We obtained similar results in the sponge *Sycon* and the cnidarian *Aglantha* (**Figs. 4A, 2S**-supplement). Although we do not entirely exclude the presence of D-glutamate in placozoans, sponges, and cnidarians (maybe at trace amounts or in specific cells), D-aspartate is a prevail enantiomer in these three basally metazoan lineages as in Bilateria (**Fig. 4B**).

## 4. Discussion

Recent phylogenomic reconstructions suggest that Ctenophores are the earliest-branching lineage of all animals [40,43]. The second branch on the animal tree of life could be the lineage leading to sponges (or Porifera, see also [44]). The current consensus also stands that Placozoa is the sister group to the clade Cnidaria+Bilarteria [40,43,44] and **Fig. 4B**. Regardless of the proposed phylogenies, these four groups represent crucial taxon sampling to understand the origin and evolution of animal traits, the nervous system, and neurotransmitters in particular [45].

We showed that the placozoans, cnidarians, and sponges had high micromolar concentrations of D-aspartate, whereas D-glutamate was not detectable. In contrast, D-glutamate was the main enantiomer in ctenophores, with no or trace amount of D-aspartate (**Fig.4**). This situation illuminates lineage-specific diversifications in the recruitment of D-amino acids and suggests distinct signaling functions of these molecules early in the animal evolution. This hypothesis might explain the observed relatively low abundance (or reduced recruitment) of D-glutamate in neural systems of bilaterians. It could be a result of the early split and parallel the evolution of D-amino acids’ signaling systems from the common metazoan ancestor to the lineage led to ctenophores (predominantly D-glutamate) and the rest of the animals (predominantly D-aspartate), respectively.

It was shown that in ctenophores, both D- and L-glutamate induced muscle contractions and elevation of intracellular Ca^2+^ [43]. Thus, one or two enantiomers of glutamate could act as a neuromuscular transmitter(s) in ctenophores. The functions of D-aspartate in sponges, placozoans, and cnidarians are unknown, but potentially D-amino acids can be incorporated in toxins and signaling peptides as in bilaterians [46].

What would be likely sources of D-glutamate in ctenophores and D-aspartate in placozoans, sponges, and cnidarians? Dietary, bacterial/symbiotic origins and enzymatic synthesis are possible sources to be explored. At least two serine/aspartate racemases have been implicated in the synthesis of D-aspartate in bilaterians [34,35,47]. Nonetheless, the synthesis of D-glutamate is largely unresolved. D-glutamate might be produced by lineage-specific and genetically unrelated enzymes. For example, a specialized aspartate racemase(SSS) can catalyze L- to D-glutamate conversiton only in hemichordates [24].

We did not find any obvious candidate for genes encoding D-glutamate synthetic or degradation enzymes in the sequenced genomes of *Pleurobrachia* and *Mnemiopsis* (such as D-aspartate oxidase [EC1.4.3.1] or D-glutamate cyclase [EC4.2.1.48]). Yet, in ctenophores, we found genes encoding putative serine racemase/dehydrogenase, which might be candidates for future expression studies and enzymatic assays (Fig 1S, Supplement-1). Similar orthologs can also be explored as candidates for L- to D-aspartate conversion in sponges and cnidarians. There is just one putative racemase with unknown specificity in *Trichoplax* (Supplement-1). Thus, we expect the discovery of novel enzymatic and nonenzymatic pathways of D-amino acid synthesis and catabolism at the base of the animal tree of life.

Finally, we did not exclude a possibility for the presence of D-aspartate in specific cells of ctenophores and D-glutamate in sponges, placozoans, and cnidarians. Some improvements in LODs and single-cell methods could be implemented to minimize errors and expand comparative surveys[45]. However, our study suggests the early-evolutionary split in the recruitment of D-glutamate between ctenophores and the rest of the animals. And, a broader sampling would be required to test further the hypothesis outlined in **Fig. 4B**.

## Acknowledgments

This work was supported by the Human Frontiers Science Program (RGP0060/2017) and National Science Foundation (1146575,1557923,1548121,1645219) grants to L.L.M

We thank the Friday Harbor Laboratories for their support to collect animals.

## Role of authors

All authors had access to the data in the study and take responsibility for the integrity of the data and the accuracy of the data analysis. Research design, writing: LLM. Acquisition of data: all authors. Molecular data: DYR, ABK, LLM; Microchemical assays: DS, LLM; animal collections, dissections: LLM, DYR; sample preparations: LLM, DS. Analysis and interpretation: all authors.: Funding: LLM.

---

**Fig 1S-Supplement.**
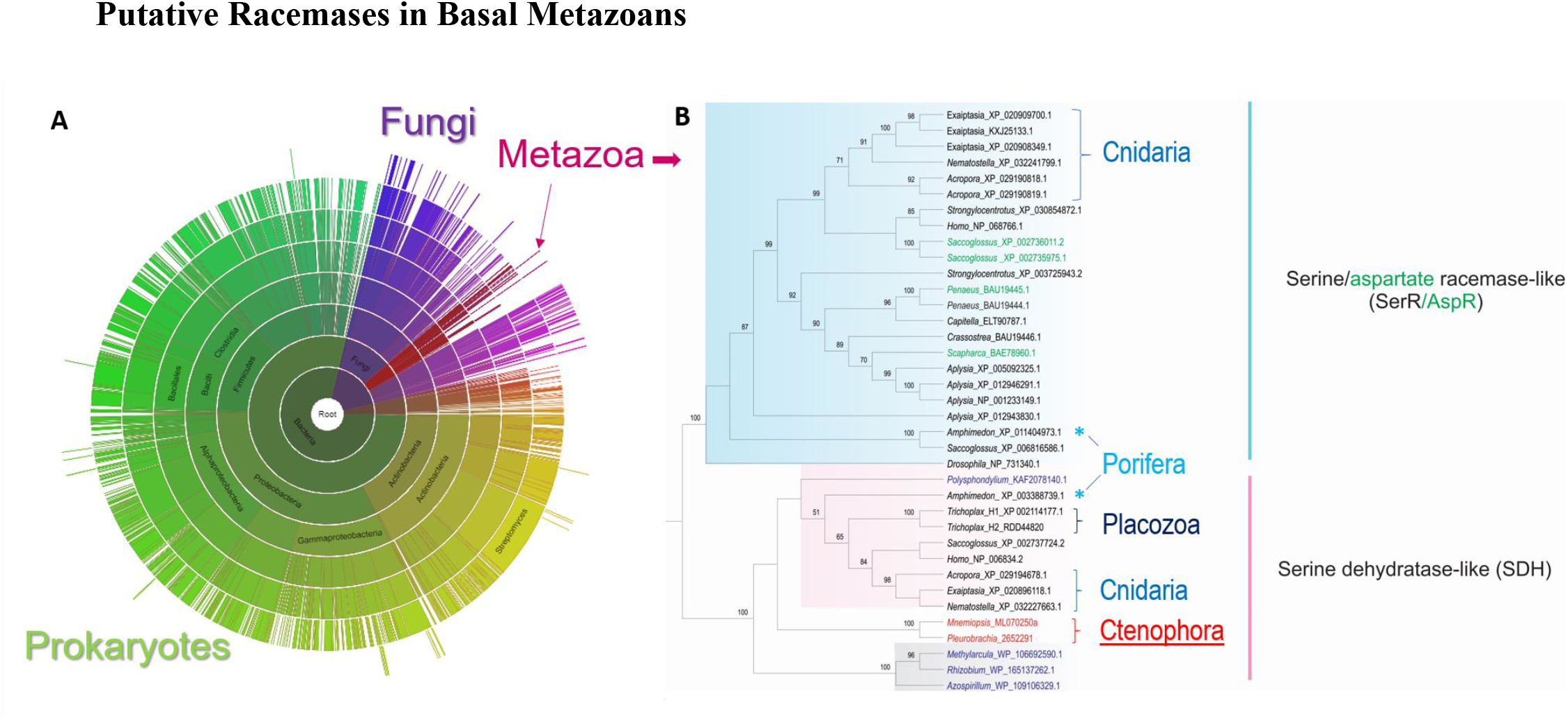
Phylogenetic tree of Serine dehydrogenase/Serine racemase orthologs in representative animal lineages.

**Fig. 2S-Supplement.**
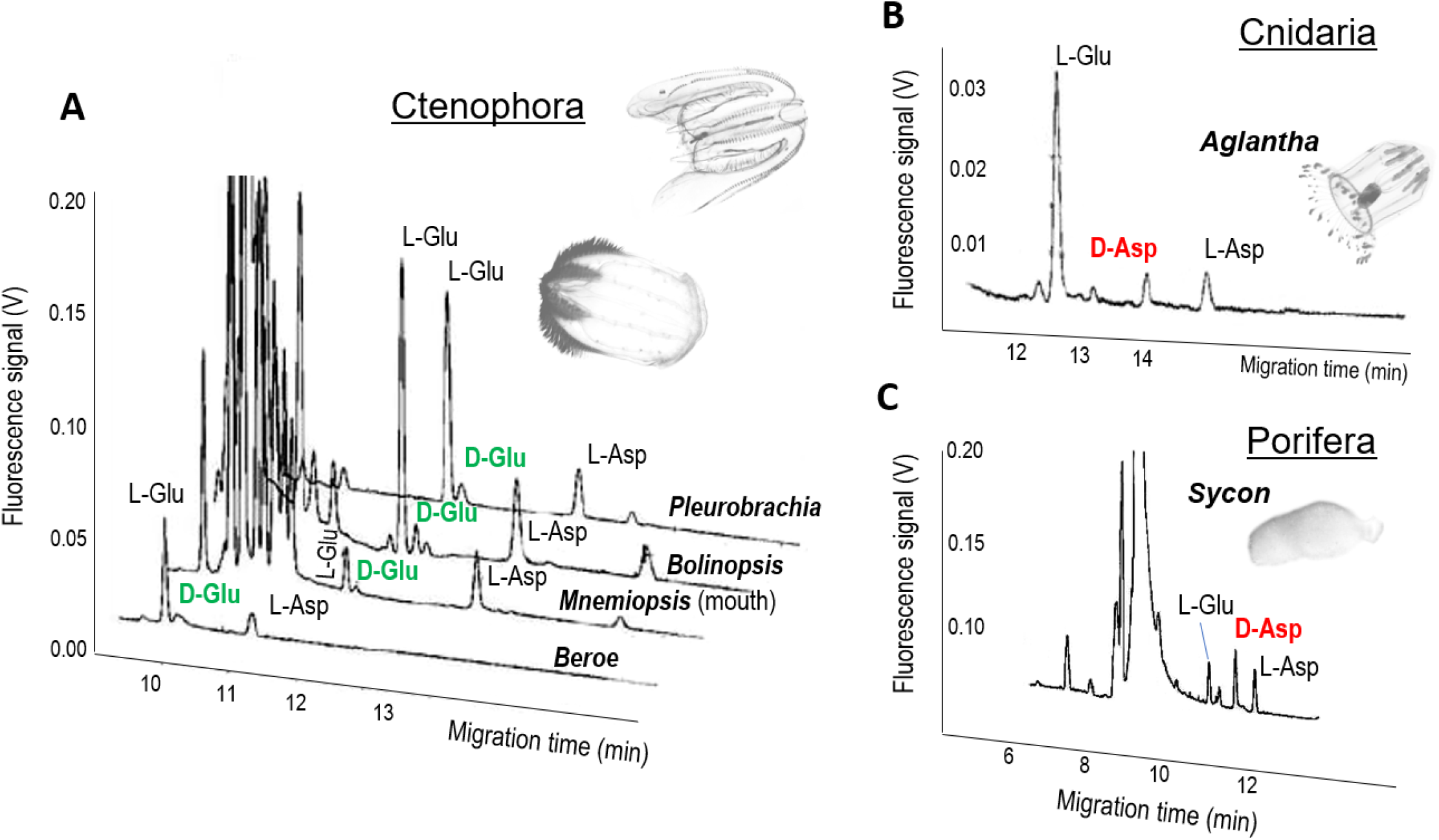
Examples of the enantiomer separations in ctenophores, cnidarians, and sponges.

**Fig. 3S-Supplement.**
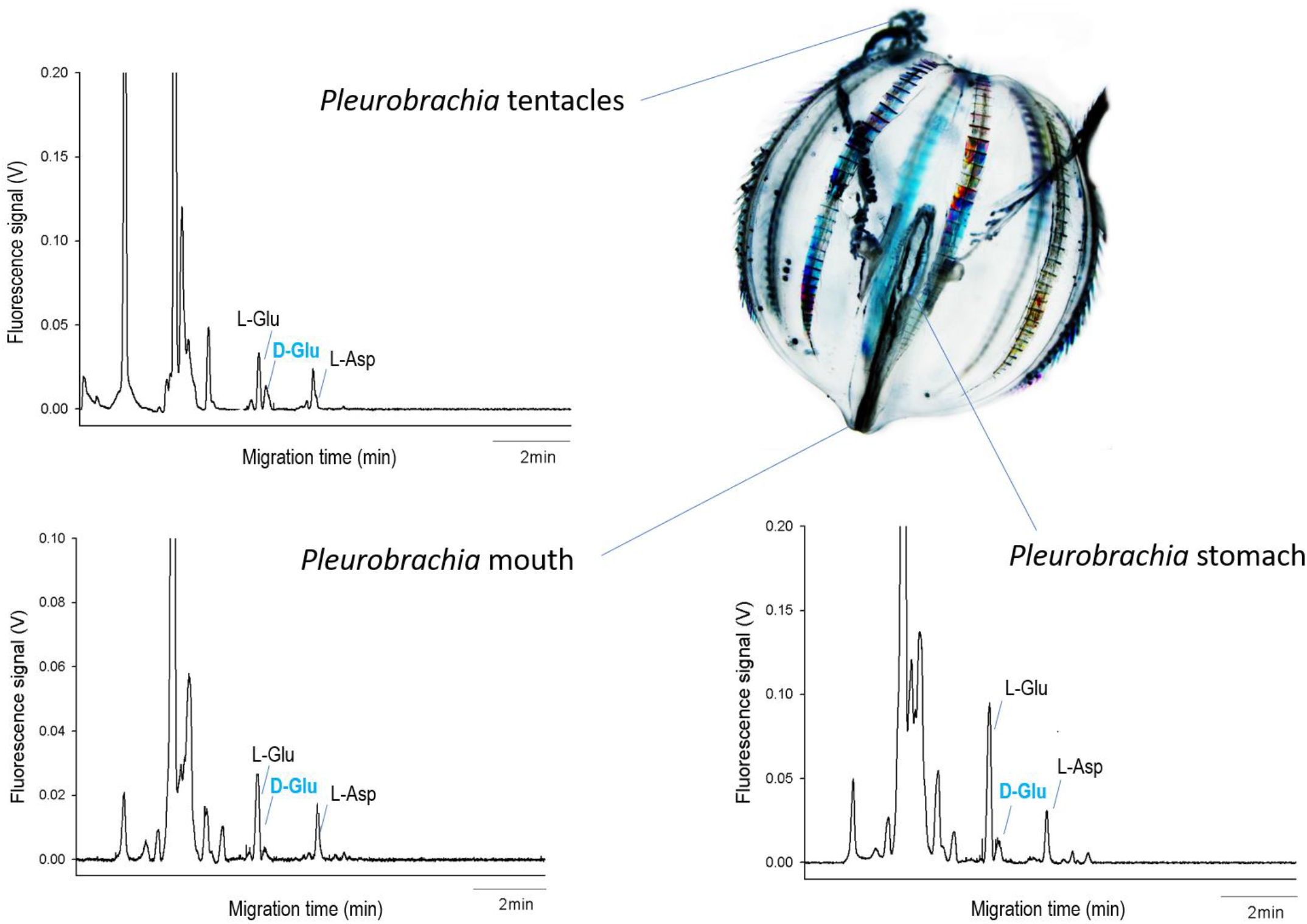
Detections of D-glutamate in different tissues of *Pleurobrachia bachei*.

## Supplement-2

Sequences used for the reconstruction of the phylogenetic tree in **Fig.1S**.

## References

[1] S.A. Fuchs, R. Berger, L.W. Klomp, T.J. de Koning, D-amino acids in the central nervous system in health and disease, Mol Genet Metab 85 (2005) 168–180. 10.1016/j.ymgme.2005.03.003.

[2] N. Fujii, T. Takata, N. Fujii, K. Aki, H. Sakaue, D-Amino acids in protein: The mirror of life as a molecular index of aging, Biochim Biophys Acta Proteins Proteom 1866 (2018) 840–847. 10.1016/j.bbapap.2018.03.001.

[3] G. Genchi, An overview on D-amino acids, Amino Acids 49 (2017) 1521–1533. 10.1007/s00726-017-2459-5.

[4] J.J. Corrigan, D-amino acids in animals, Science 164 (1969) 142–149. 10.1126/science.164.3876.142.

[5] H. Rosenberg, A.H. Ennor, The occurrence of free D-serine in the earthworm, Biochem J 79 (1961) 424–428. 10.1042/bj0790424.

[6] H. Rosenberg, A.H. Ennor, Occurrence of free D-serine in the earthworm, Nature 187 (1960) 617–618. 10.1038/187617a0.

[7] Z. Feng, B. Xu, Inspiration from the mirror: D-amino acid containing peptides in biomedical approaches, Biomol Concepts 7 (2016) 179–187. 10.1515/bmc-2015-0035.

[8] J. Sasabe, M. Suzuki, Emerging Role of D-Amino Acid Metabolism in the Innate Defense, Front Microbiol 9 (2018) 933. 10.3389/fmicb.2018.00933.

[9] E.H. Man, M.E. Sandhouse, J. Burg, G.H. Fisher, Accumulation of D-aspartic acid with age in the human brain, Science 220 (1983) 1407–1408. 10.1126/science.6857259.

[10] D.S. Dunlop, A. Neidle, D. McHale, D.M. Dunlop, A. Lajtha, The presence of free D-aspartic acid in rodents and man, Biochem Biophys Res Commun 141 (1986) 27–32. 10.1016/s0006-291x(86)80329-1.

[11] H. Wolosker, E. Dumin, L. Balan, V.N. Foltyn, D-amino acids in the brain: D-serine in neurotransmission and neurodegeneration, FEBS J 275 (2008) 3514–3526. 10.1111/j.1742-4658.2008.06515.x.

[12] A.K. Mustafa, P.M. Kim, S.H. Snyder, D-Serine as a putative glial neurotransmitter, Neuron Glia Biol 1 (2004) 275–281. 10.1017/S1740925X05000141.

[13] P. Fossat, F.R. Turpin, S. Sacchi, J. Dulong, T. Shi, J.M. Rivet, J.V. Sweedler, L. Pollegioni, M.J. Millan, S.H. Oliet, J.P. Mothet, Glial D-serine gates NMDA receptors at excitatory synapses in prefrontal cortex, Cereb Cortex 22 (2012) 595–606. 10.1093/cercor/bhr130.

[14] N. Ota, T. Shi, J.V. Sweedler, D-Aspartate acts as a signaling molecule in nervous and neuroendocrine systems, Amino Acids 43 (2012) 1873–1886. 10.1007/s00726-012-1364-1.

[15] A. Mangas, R. Covenas, D. Bodet, M. Geffard, L.A. Aguilar, J. Yajeya, Immunocytochemical visualization of D-glutamate in the rat brain, Neuroscience 144 (2007) 654–664. 10.1016/j.neuroscience.2006.09.045.

[16] R. Covenas, A. Mangas, M.L. Sanchez, D. Cadena, M. Husson, M. Geffard, Generation of specific antisera directed against D-amino acids: focus on the neuroanatomical distribution of D-glutamate and other D-amino acids, Folia Histochem Cytobiol 55 (2017) 177–189. 10.5603/FHC.a2017.0023.

[17] C.A. Weatherly, S. Du, C. Parpia, P.T. Santos, A.L. Hartman, D.W. Armstrong, d-Amino Acid Levels in Perfused Mouse Brain Tissue and Blood: A Comparative Study, ACS Chem Neurosci 8 (2017) 1251–1261. 10.1021/acschemneuro.6b00398.

[18] A.V. Patel, T. Kawai, L. Wang, S.S. Rubakhin, J.V. Sweedler, Chiral Measurement of Aspartate and Glutamate in Single Neurons by Large-Volume Sample Stacking Capillary Electrophoresis, Anal Chem 89 (2017) 12375–12382. 10.1021/acs.analchem.7b03435.

[19] P. Spinelli, E.R. Brown, G. Ferrandino, M. Branno, P.G. Montarolo, E. D’Aniello, R.K. Rastogi, B. D’Aniello, G.C. Baccari, G. Fisher, A. D’Aniello, D-aspartic acid in the nervous system of *Aplysia limacina*: possible role in neurotransmission, J Cell Physiol 206 (2006) 672–681. 10.1002/jcp.20513.

[20] N. Yoshikawa, W. Ashida, K. Hamase, H. Abe, HPLC determination of the distribution of D-amino acids and effects of ecdysis on alanine racemase activity in kuruma prawn *Marsupenaeus japonicus*, J Chromatogr B Analyt Technol Biomed Life Sci 879 (2011) 3283–3288. 10.1016/j.jchromb.2011.04.026.

[21] K. Sekimizu, J. Larranaga, H. Hamamoto, M. Sekine, T. Furuchi, M. Katane, H. Homma, N. Matsuki, D-Glutamic acid-induced muscle contraction in the silkworm, *Bombyx mori*, J Biochem 137 (2005) 199–203. 10.1093/jb/mvi019.

[22] J.J. Corrigan, N.G. Srinivasan, The occurrence of certain D-amino acids in insects, Biochemistry 5 (1966) 1185–1190. 10.1021/bi00868a010.

[23] K. Shibata, N. Sugaya, Y. Kuboki, H. Matsuda, K. Abe, S. Takahashi, Y. Kera, Aspartate racemase and D-aspartate in starfish; possible involvement in testicular maturation, Biosci Biotechnol Biochem 84 (2020) 95–102. 10.1080/09168451.2019.1660614.

[24] K. Uda, N. Ishizuka, Y. Edashige, A. Kikuchi, A.D. Radkov, L.A. Moe, Cloning and characterization of a novel aspartate/glutamate racemase from the acorn worm *Saccoglossus kowalevskii*, Comp Biochem Physiol B Biochem Mol Biol 232 (2019) 87–92. 10.1016/j.cbpb.2019.03.006.

[25] S. D’Aniello, P. Spinelli, G. Ferrandino, K. Peterson, M. Tsesarskia, G. Fisher, A. D’Aniello, Cephalopod vision involves dicarboxylic amino acids: D-aspartate, L-aspartate and L-glutamate, Biochem J 386 (2005) 331–340. 10.1042/BJ20041070.

[26] B. De Marianis, A. Giuditta, Separation of nuclei with different DNA content from the suboesophageal lobe of octopus brain, Brain Res 154 (1978) 134–136. 10.1016/0006-8993(78)91059-4.

[27] S. D’Aniello, I. Somorjai, J. Garcia-Fernandez, E. Topo, A. D’Aniello, D-Aspartic acid is a novel endogenous neurotransmitter, FASEB J 25 (2011) 1014-1027. 10.1096/fj.10-168492.

[28] Y.M. Liu, M. Schneider, C.M. Sticha, T. Toyooka, J.V. Sweedler, Separation of amino acid and peptide stereoisomers by nonionic micelle-mediated capillary electrophoresis after chiral derivatization, J Chromatogr A 800 (1998) 345–354. 10.1016/s0021-9673(97)01137-0.

[29] H. Miao, S.S. Rubakhin, J.V. Sweedler, Subcellular analysis of D-aspartate, Anal Chem 77 (2005) 7190–7194. 10.1021/ac0511694.

[30] H. Miao, S.S. Rubakhin, C.R. Scanlan, L. Wang, J.V. Sweedler, D-Aspartate as a putative cell-cell signaling molecule in the Aplysia californica central nervous system, J Neurochem 97 (2006) 595–606. 10.1111/j.1471-4159.2006.03791.x.

[31] H. Miao, S.S. Rubakhin, J.V. Sweedler, Confirmation of peak assignments in capillary electrophoresis using immunoprecipitation. Application to D-aspartate measurements in neurons, J Chromatogr A 1106 (2006) 56–60. 10.1016/j.chroma.2005.09.037.

[32] C. Scanlan, T. Shi, N.G. Hatcher, S.S. Rubakhin, J.V. Sweedler, Synthesis, accumulation, and release of d-aspartate in the Aplysia californica CNS, J Neurochem 115 (2010) 1234–1244. 10.1111/j.1471-4159.2010.07020.x.

[33] Y. Saitoh, M. Katane, T. Kawata, K. Maeda, M. Sekine, T. Furuchi, H. Kobuna, T. Sakamoto, T. Inoue, H. Arai, Y. Nakagawa, H. Homma, Spatiotemporal localization of D-amino acid oxidase and D-aspartate oxidases during development in *Caenorhabditis elegans*, Mol Cell Biol 32 (2012) 1967–1983. 10.1128/MCB.06513-11.

[34] K. Uda, K. Abe, Y. Dehara, K. Mizobata, Y. Edashige, R. Nishimura, A.D. Radkov, L.A. Moe, Triple serine loop region regulates the aspartate racemase activity of the serine/aspartate racemase family, Amino Acids 49 (2017) 1743–1754. 10.1007/s00726-017-2472-8.

[35] K. Uda, K. Abe, Y. Dehara, K. Mizobata, N. Sogawa, Y. Akagi, M. Saigan, A.D. Radkov, L.A. Moe, Distribution and evolution of the serine/aspartate racemase family in invertebrates, Amino Acids 48 (2016) 387–402. 10.1007/s00726-015-2092-0.

[36] D.Y. Romanova, A. Heyland, D. Sohn, A.B. Kohn, D. Fasshauer, F. Varoqueaux, L.L. Moroz, Glycine as a signaling molecule and chemoattractant in *Trichoplax* (Placozoa): insights into the early evolution of neurotransmitters, NeuroReport 31 (2020) 490–497. 10.1097/WNR.0000000000001436

[37] P.D. Floyd, L.L. Moroz, R. Gillette, J.V. Sweedler, Capillary electrophoresis analysis of nitric oxide synthase related metabolites in single identified neurons, Anal Chem 70 (1998) 2243–2247. 10.1021/ac9713013.

[38] L.L. Moroz, R.L. Dahlgren, D. Boudko, J.V. Sweedler, P. Lovell, Direct single cell determination of nitric oxide synthase related metabolites in identified nitrergic neurons, J Inorg Biochem 99 (2005) 929–939. 10.1016/j.jinorgbio.2005.01.013.

[39] Z. Juvancz, R.B. Kendrovics, R. Ivanyi, L. Szente, The role of cyclodextrins in chiral capillary electrophoresis, Electrophoresis 29 (2008) 1701–1712.

[40] N.V. Whelan, K.M. Kocot, T.P. Moroz, K. Mukherjee, P. Williams, G. Paulay, L.L. Moroz, K.M. Halanych, Ctenophore relationships and their placement as the sister group to all other animals, Nat Ecol Evol 1 (2017) 1737–1746. 10.1038/s41559-017-0331-3.

[41] T.P. Norekian, L.L. Moroz, Neuromuscular organization of the Ctenophore *Pleurobrachia bachei*, J Comp Neurol 527 (2019) 406–436. 10.1002/cne.24546.

[42] T.P. Norekian, L.L. Moroz, Neural system and receptor diversity in the ctenophore *Beroe abyssicola*, J Comp Neurol 527 (2019) 1986–2008. 10.1002/cne.24633.

[43] L.L. Moroz, K.M. Kocot, M.R. Citarella, S. Dosung, T.P. Norekian, I.S. Povolotskaya, A.P. Grigorenko, C. Dailey, E. Berezikov, K.M. Buckley, A. Ptitsyn, D. Reshetov, K. Mukherjee, T.P. Moroz, Y. Bobkova, F. Yu, V.V. Kapitonov, J. Jurka, Y.V. Bobkov, J.J. Swore, D.O. Girardo, A. Fodor, F. Gusev, R. Sanford, R. Bruders, E. Kittler, C.E. Mills, J.P. Rast, R. Derelle, V.V. Solovyev, F.A. Kondrashov, B.J. Swalla, J.V. Sweedler, E.I. Rogaev, K.M. Halanych, A.B. Kohn, The ctenophore genome and the evolutionary origins of neural systems, Nature 510 (2014) 109–114. 10.1038/nature13400.

[44] C.E. Laumer, R. Fernandez, S. Lemer, D. Combosch, K.M. Kocot, A. Riesgo, S.C.S. Andrade, W. Sterrer, M.V. Sorensen, G. Giribet, Revisiting metazoan phylogeny with genomic sampling of all phyla, Proc Biol Sci 286 (2019) 20190831. 10.1098/rspb.2019.0831.

[45] L.L. Moroz, NeuroSystematics and Periodic System of Neurons: Model vs Reference Species at Single-Cell Resolution, ACS Chem Neurosci 9 (2018) 1884–1903. 10.1021/acschemneuro.8b00100.

[46] C. Ollivaux, D. Soyez, J.Y. Toullec, Biogenesis of D-amino acid containing peptides/proteins: where, when and how?, J Pept Sci 20 (2014) 595–612. 10.1002/psc.2637.

[47] T. Ito, M. Hayashida, S. Kobayashi, N. Muto, A. Hayashi, T. Yoshimura, H. Mori, Serine racemase is involved in d-aspartate biosynthesis, J Biochem 160 (2016) 345–353. 10.1093/jb/mvw043.

